# Standardization of Data Analysis for RT-QuIC-based Detection of Chronic Wasting Disease

**DOI:** 10.1101/2022.11.17.516950

**Authors:** Gage R. Rowden, Catalina Picasso-Risso, Manci Li, Marc D. Schwabenlander, Tiffany Wolf, Peter Larsen

## Abstract

Chronic wasting disease (CWD) is a disease affecting cervids and is caused by prions accumulating as pathogenic fibrils in lymphoid tissue and the central nervous system. Approaches for detecting CWD prions historically relied on antibody-based assays. However, recent advancements in protein amplification technology provided the foundation for a new class of CWD diagnostic tools. In particular, real-time quaking-induced conversion (RT-QuIC) has rapidly become a feasible option for CWD diagnosis. Despite its increased usage for CWD-focused research, there lacks consensus regarding the interpretation of RT-QuIC data for diagnostic purposes. It is imperative then to identify a standardized and replicable method for determining CWD status from RT-QuIC data. Here, we assessed variables that could impact RT-QuIC results and explored the use of maxpoint ratios (maximumRFU/backgroundRFU) to improve the consistency of RT-QuIC analysis. We examined a variety of statistical analyses to retrospectively analyze CWD status based on RT-QuIC and ELISA results from 668 white-tailed deer lymph nodes. Our results revealed an MPR threshold of 2.0 for determining the rate of amyloid formation, and MPR analysis showed excellent agreement with independent ELISA results. These findings suggest that the use of MPR is a statistically viable option for normalizing between RT-QuIC experiments and defining CWD status.

## Introduction

The diagnosis of prion diseases has historically relied on immunoassays such as enzyme-linked immunosorbent assay (ELISA) and immunohistochemistry (IHC). However, recent advancements in protein amplification assays such as real-time quaking induced conversion (RT-QuIC) have emerged as powerful tools for detecting a variety of transmissible spongiform encephalopathies (TSEs) and other proteopathies including chronic wasting disease (CWD), scrapie, bovine spongiform encephalopathy (BSE), Creutzfeldt-Jakob disease (CJD), Alzheimer’s disease, and Parkinson’s disease [1, 3, 8, 14, 30, 36, 42, 43]. CWD is caused by the misfolding of prion protein (PrP*^CWD^*) and subsequent recruitment of native prion protein (PrPC) into pathological amyloid fibrils, thus causing a contagious and unconditionally fatal disease [46, 47]. The disease was first described in captive mule deer in Colorado in 1967 and has since been identified across the United States and Canada, Scandinavia, and South Korea [39, 52, 57]. Because of the severity and continued spread of the disease, diagnostic methods must evolve to meet the growing need for fast and accurate testing. RT-QuIC quantifies amyloid fibril formation via relative fluorescence (RFU) using thioflavin T (ThT) as a fluorescent marker and has shown similar-to-improved accuracy when compared to IHC and ELISA [24, 45]. Moreover, the United States Department of Agriculture Animal and Plant Health Inspection Service (USDA APHIS) is currently validating RT-QuIC for the routine detection of CWD in farmed cervids (Tracy Nichols; personal communication).

Despite the growing interest in RT-QuIC-based diagnostics, there is currently no consensus regarding the interpretation of the assay’s output. Most authors determine CWD positivity by setting a predefined number of standard deviations (e.g., between 3–30) above the average background ThT fluorescence per plate and defining that fluorescent value as a positivity threshold (T*_stdev_*). A rate of amyloid formation (RAF) is then calculated as the inverse of the time that the reaction took to pass T*_stdev_*. Finally, a positive sample is determined arbitrarily when some minimum percentage (generally between 25–67%) of replicates passes T*_stdev_* [2–4, 6–13, 15–23, 25–35, 37,38, 40–44, 48–51, 53–55, 58,59] (Table 1).

**Table 1.** A compilation of the various methods used in publications for determining an RT-QuIC-positive sample when multiple replicates are processed. Positivity is typically assessed by a predetermined number of replicates crossing a study-specific threshold that is variably defined between 3–30 standard deviations from the initial mean relative fluorescent units (RFU) of the entire plate. CWD: chronic wasting disease; CJD: Creutzfeldt-Jakob disease; PD: Parkinson’s disease; AD: Alzheimer’s disease; BSE: bovine spongiform encephalopathy; FFI: fatal familial insomnia; Sc: scrapie; stdev: standard deviations

| Threshold used | Positivity Determination | Disease | Reference |
| --- | --- | --- | --- |
| 20 stdev | Mann-Whitney u-test | AD | [54] |
| 10 stdev | 50% of replicates | BSE | [11] [27] |
| 10 stdev | 25% of replicates | BSE | [30] |
| 20 stdev | 50% of replicates | CJD | [44] |
| 10 stdev+10% | 50% of replicates | CJD | [43] |
| 10 stdev | 50% of replicates | CJD | [12] [2] [4] [13] [41] |
| 5 stdev | 50% of replicates | CJD | [38] |
| 5 stdev | 40% of replicates | CJD | [37] |
| 3 stdev | 50% of replicates | CJD | [59] [33] |
| >10,000 RFU | 50% of replicates | CJD | [7] |
| 10 stdev | Mann-Whitney u-test | CWD | [31] [51] |
| 10 stdev | 2/3 of replicates | CWD | [16] |
| 10 stdev | 50% of replicates | CWD | [17] [29] |
| 10 stdev | 25% of replicates | CWD | [28] |
| 10 stdev | 1/3 of replicates | CWD | [18] [29] |
| 10 stdev | Avg of reps $>T_{stdev}$ | CWD | [19] [32] |
| 5 stdev | Mann-Whitney or Wilcoxon signed rank | CWD | [35] |
| 5 stdev | Mann-Whitney u-test | CWD | [55] [40] [34] |
| 5 stdev | Fisher’s exact test | CWD | [23] |
| 5 stdev | Unpaired t-test | CWD | [25] |
| 5 stdev | One sample t-test | CWD | [26] |
| 5 stdev | 50% of replicates | CWD | [6] |
| 5 stdev | Avg of reps $>T_{stdev}$ | CWD | [22] [15] |
| 4 stdev | 50% of replicates | CWD | [10] |
| max signal of pos ctrl | 1/3 of replicates | CWD | [48] |
| 5 stdev | Mann-Whitney or Wilcoxon signed rank | CWD; BSE | [9] |
| 5 stdev | Avg of reps $>T_{stdev}$ | CWD; BSE | [53] |
| 3 stdev | 50% of replicates | FFI | [58] |
| 30 stdev | 50% of replicates | PD | [49] |
| 10 stdev+10% | 50% of replicates | PD | [21] [3] |
| >120 RFU | 50% of replicates | PD | [50] |
| 10 stdev | 50% of replicates | Sc | [8] |

Additionally, some researchers incorporate statistical analyses, such as Mann-Whitney, Wilcoxon signed-rank, or Fisher’s exact test which statistically compares the RAF replicates of unknown-CWD-status samples to a control sample [9, 31, 35, 40, 51, 54, 55]. Although RAFs can provide a quantitative assessment of relative PrP*^CWD^* load, they are not statistically informative when compared to a negative control, and CWD determinations from their analyses are ultimately susceptible to researcher preference. This challenges standardization among laboratories and the external validity of results. Given the increased usage of RT-QuIC for CWD research and surveillance, a robust assessment of the analytical process to define disease status with RT-QuIC data is needed.

Here, we describe a method for determining CWD status of a given sample based on what we have coined the maxpoint ratio (MPR) [60]. This method was adapted from Cheng et al. [5, 56] and corrects replicates for background fluorescence and normalizes data output between experiments. CWD-positivity determinations using MPR are based on a statistical analysis against the MPR values of a known negative control rather than deciding beforehand the necessary number of replicates needed to cross a threshold. Nevertheless, the RAF of a reaction is still an important kinetic measurement which gives an idea of relative prion load in a sample, so a constant threshold (T*_MPR_*) was determined at twice the background fluorescence for RAF calculations rather than assigning independent thresholds (T*_stdev_*) per plate. With the use of this proposed method we aim to normalize results across RT-QuIC experiments, thus standardizing RT-QuIC diagnostic results across laboratories and improving CWD control programs.

## Materials and Methods

### Literature Review

To identify the methods used by researchers to determine TSE status from RT-QuIC data, we conducted a PubMed search (keywords: RT-QuIC and “real-time quaking-induced conversion”) for articles that performed RT-QuIC between 2012 and 2021. Studies were selected if the researchers both performed RT-QuIC and implemented a rubric for determining TSE status. Studies were not vetted for any particular prion disease or proteopathy.

### Source Population and Sample Processing

In accordance with the culling efforts of the Minnesota Department of Natural Resources (DNR), 668 lymph node samples from 533 wild white-tailed deer (WTD) were sampled from CWD endemic areas in southwestern Minnesota [51]. Following the DNR’s CWD surveillance program mentioned above, a pooled homogenate of three sub-samples for each animal’s RPLNs were submitted for independent screening by ELISA to the Colorado State University Veterinary Diagnostic Laboratory (CSU VDL, Fort Collins, Colorado).

Additionally, parotid (PLN; n=515), medial retropharyngeal (RPLN, n=17), submandibular (SMLN; 63), popliteal (PPLN; n=1), prescapular (PSLN; n=10), and palatine tonsil (TLN; n=62) lymph nodes were collected and stored at −20 °C. Approximately 100 mg of lymph node tissue were homogenized in 1X PBS to a concentration of 10% (w:v) using a BeadBug™ homogenizer (Benchmark Scientific, Sayreville New Jersey, USA) and 1.5mm zirconium beads (Millipore Sigma, Burlington, Massachusetts, USA) for 90 seconds at the maximum setting. Homogenates were stored at −80 °C until tested, at which point they were diluted 100-fold (10-3 final dilution) in dilution buffer (1X PBS, 0.1% [w:v] sodium dodecyl sulfate, 1X N_2_ supplement [Life Technologies Corporation, Carlsbad, California, USA]) and were vortexed [51]. RT-QuIC was performed at the Minnesota Center for Prion Research and Outreach (MNPRO; see below).

Furthermore, a subset of PLNs, SMLNs, and TLNs from 60 WTD were selected and submitted to the CSU VDL for blind testing with ELISA and by RT-QuIC at MNPRO to determine concordance between the two assays. ELISAs were performed with the Bio-Rad TeSeE Short Assay Protocol (SAP) Combo Kit (BioRad Laboratories Inc., Hercules, CA, USA). Importantly, while the Bio-Rad ELISA is used as the standard screening assay for CWD, it is only currently validated for RPLNs and obex.

### RT-QuIC Methods and recPrP Preparation

Expression and purification of recombinant prion protein (recPrP; hamster, amino acids 90-231) and RT-QuIC assays were performed following Schwabenlander et al. [51] Briefly, 98 *μ**L*** of the reaction master mix and 2 μL of the 10^-3^ homogenate in dilution buffer were added to each well of a 96-well plate. The final tissue concentration was 0.002% (w:v). Reactions were performed in quadruplicate for unknown samples (n=668), and six replicates for controls (n=2). All experiments were run using four BMG FluoStar® plate readers (BMG Labtech, Ortenberg, Germany). Four different batches of recPrP substrate were used in the span of this study and were produced at The University of Minnesota Biotechnology Resource Center.

### Maxpoint Ratio and Rate of Amyloid Formation Calculations

MPRs and their associated RAF values were adapted from the algorithm of Cheng, et al. [5] MPRs were calculated by dividing the maximum RFU value obtained within 48 hours by the background RFU value for each individual replicate (i.e. MPR = maxRFU/backgroundRFU). MPRs from each replicate were plotted using GraphPad Prism version 9.0.1 (GraphPad Software, San Diego, California, USA) to determine any observable distribution patterns (Figure 1A). A constant T*_MPR_* of two was chosen as a convenient yet empirical cutoff for determining RAF. A formal assessment of the performance of T*_MPR_* was determined by ROC analysis by comparing individual MPR values to tissue-matched ELISA results (Figure 3). Cutoff values were listed in descending order of Youden’s index (*J* = sensitivity + specificity - 1). RAFs were computed retrospectively by taking the reciprocal of the time in seconds needed to cross T*_MPR_*. If a replicate did not cross T*_MPR_* within 48 hours, it was assigned an RAF of zero (see Supplementary Figure 1).

**Figure 1.**
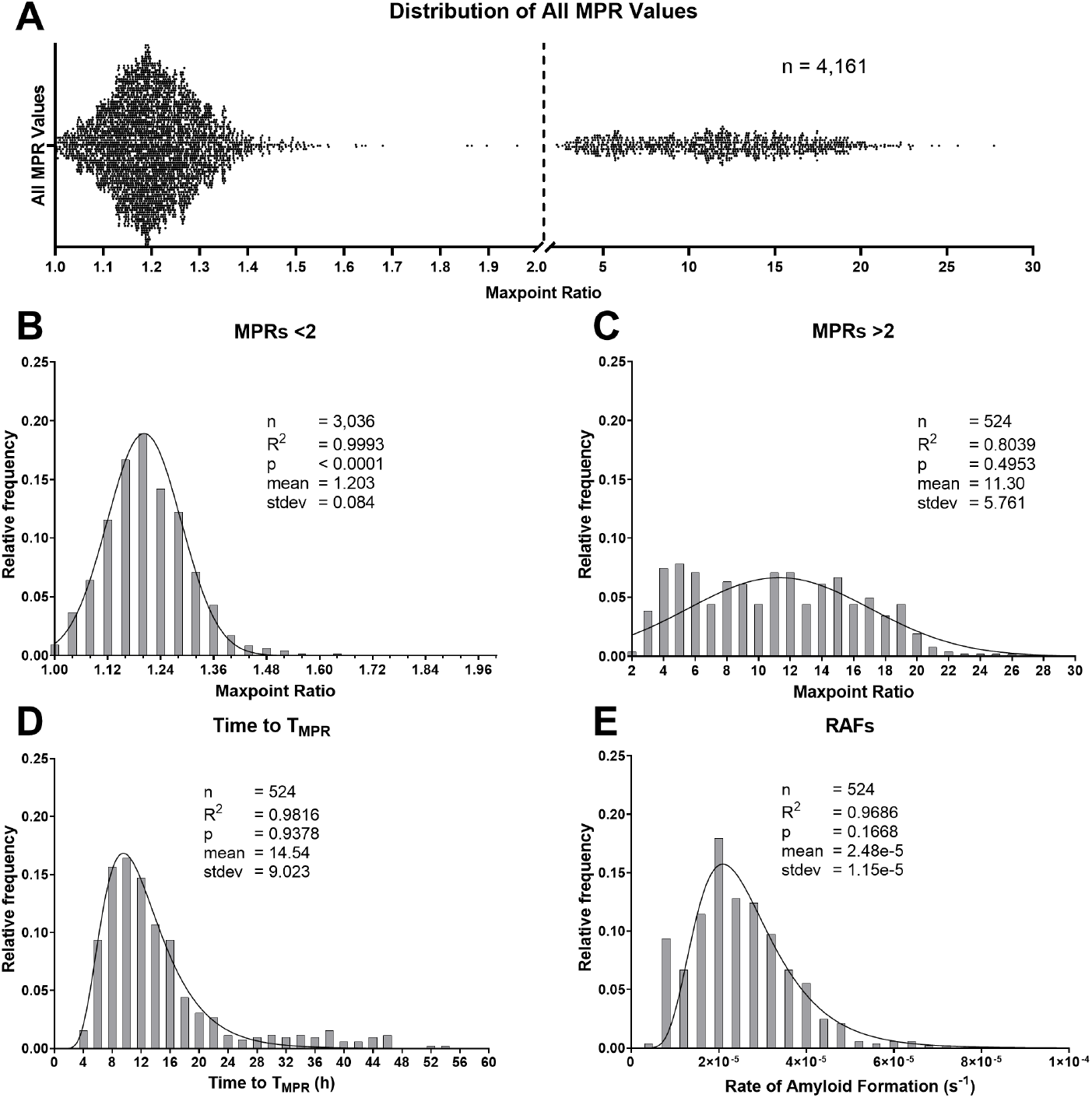
All maxpoint ratios (MPR) showing two distinct distributions separated by the dotted line at 2.0. **B.** The frequency distribution of replicates with MPR values <2.0. It does not include negative controls (n=285). These MPRs exhibited a Gaussian distribution, and the low p-value indicated that the distribution was skewed in the positive direction. **C** Distribution of the MPRs >2.0 minus the control (n=316) MPRs. The data were distributed normally although 20% of the variance was unexplained by the regression model. **D,E**: Time-to-threshold values and their corresponding rates of amyloid formation (RAF) were distributed in a lognormal pattern. stdev: standard deviation

### Statistical Analysis

Data curation and management were performed in R software v4.1.0 (R Core Team 2021), and GraphPad Prism version 9.0.1 (GraphPad Software, San Diego, CA, USA) was used for all statistical analyses. Relative frequency distributions and density plots for MPR values were constructed for both unknown (n=3,560) and control (n=601) samples. For those replicates which crossed T*_MPR_*, frequency distributions and density plots were constructed for the TtT and RAFs for unknown (n=524) and control (n=316) replicates. R^2^ values and p-values were calculated to diagnose the regressions (D’Agostino-Pearson omnibus [K2] normality of residuals test). A low p-value indicates that the distribution is skewed, and an R^2^ near 1 suggests that the model explains closer to 100% of the data variance. The lower MPR distribution was further divided based on three tissue types: PLN (n=2,594), SMLN (n=217), or TLN (n=427). Due to insufficient data points, RPLN (n=17), PPLN (n=30), and PSLN (n=16) replicates were excluded from this analysis. The differences in tissue-specific MPR distributions were compared using a Brown-Forsythe and Welch ANOVA, and a Dunnett’s T3 multiple comparisons test.

Control sample MPRs and RAFs were compiled to identify potential variability introduced by a particular microplate reader and/or recPrP production batch used for a given experiment, and outliers were eliminated as mentioned above. A Brown-Forsythe and Welch ANOVA and a Dunnett’s T3 multiple comparisons test were used to determine significance between readers or batches.

RAF values greater than zero (n=850) were plotted against their corresponding MPR values to determine the relationship between conversion efficiency and ThT fluorescent intensity. Spearman’s rank correlation test was performed to determine correlation between the individual replicate measurements.

CWD diagnosis using the MPR method was performed by comparing four MPR replicates to six replicates of each experiment’s corresponding negative control using a one-way ANOVA and a Dunnett’s multiple comparisons test (Figure 7B, Supplementary Figure 1). A sample was considered positive when p<0.05.

RT-QuIC results from the MPR method were compared to ELISA results when available. 420 replicates from 64 samples were analyzed using Cohen’s kappa test to determine the reliability between ELISA and RT-QuIC results using the MPR method.

## Results

### Literature Review

Literature review conducted in PubMed identified 46 peer-reviewed publications fitting our search criteria (see Table 1) (keywords: RT-QuIC and “real-time quaking-induced conversion”; between years 2012 and 2021). Upon inspection of the methods used to analyze RT-QuIC data, we discerned a common theme for defining a positivity threshold (T*_stdev_*). Forty-three studies averaged the initial fluorescence of every well on a reaction plate and defined T*_stdev_* as some number of standard deviations (between 3–30) above that average. One study [48] defined T*_stdev_* as 33% of the maximum fluorescence of the strongest positive control. Another [7] defined T*_stdev_* as 10,000 relative fluorescent units (RFU) while another [50] defined it as 120 RFU. Once T*_stdev_* was determined, three approaches were typically taken to determine positivity for a particular proteopathy: 1) some number or percentage of replicates crossing T*_stdev_*(n=30), 2) comparison to a negative control using a statistical test (n=11), or 3) the average RFU of a sample’s replicates was above T*_stdev_* (n=5).

### Maxpoint Ratio, Rate of Amyloid Formation, and Time-to-Threshold Distributions

MPR values of 4,161 individual RT-QuIC replicates from 46 reaction plates were compiled, and two distinct distributions were observed (Figure 1A). Once the distributions were determined, all control replicates were removed to account for any bias for those samples. MPRs <2.0 comprised the entirety of the lower distribution. Given this observation, we established a T*_MPR_* at an MPR value of 2.0 for calculating RAF. A density curve was fitted to the lower MPR distribution (n=3,036; R^2^: 0.9993; p<0.0001; Figure 1B). Positive MPRs distributed normally, although approximately 20% of the variance was unexplained by the model (n=524; R^2^: 0.8039; p: 0.4953; Figure 1C). However, those replicates’ time-to-threshold (TtT) (R^2^: 0.9816; p: 0.9378) and RAF values (R^2^: 0.9686; p: 0.1668) are both well-supported by their lognormal regression models (Figure 1D and 1E). The same analyses were performed on the isolated control values (Figure 2B). The pattern for negative control MPRs matched that of the lower distribution mentioned above (n=285; R^2^: 0.9993; p: 0.0513; Figure 2A). Positive control MPRs distributed normally, but approximately 10% of the variance was unexplained by the regression (n=316; R^2^: 0.8985; p: 0.2137; Figure 2C). However, the lognormal regression model for positive control RAFs was an excellent fit (n=316; R^2^: 0.9912; p: 0.5860; Figure 2D). The low p-values for both Figure 1B and Figure 2A indicate that these distributions are positively skewed.

**Figure 2.**
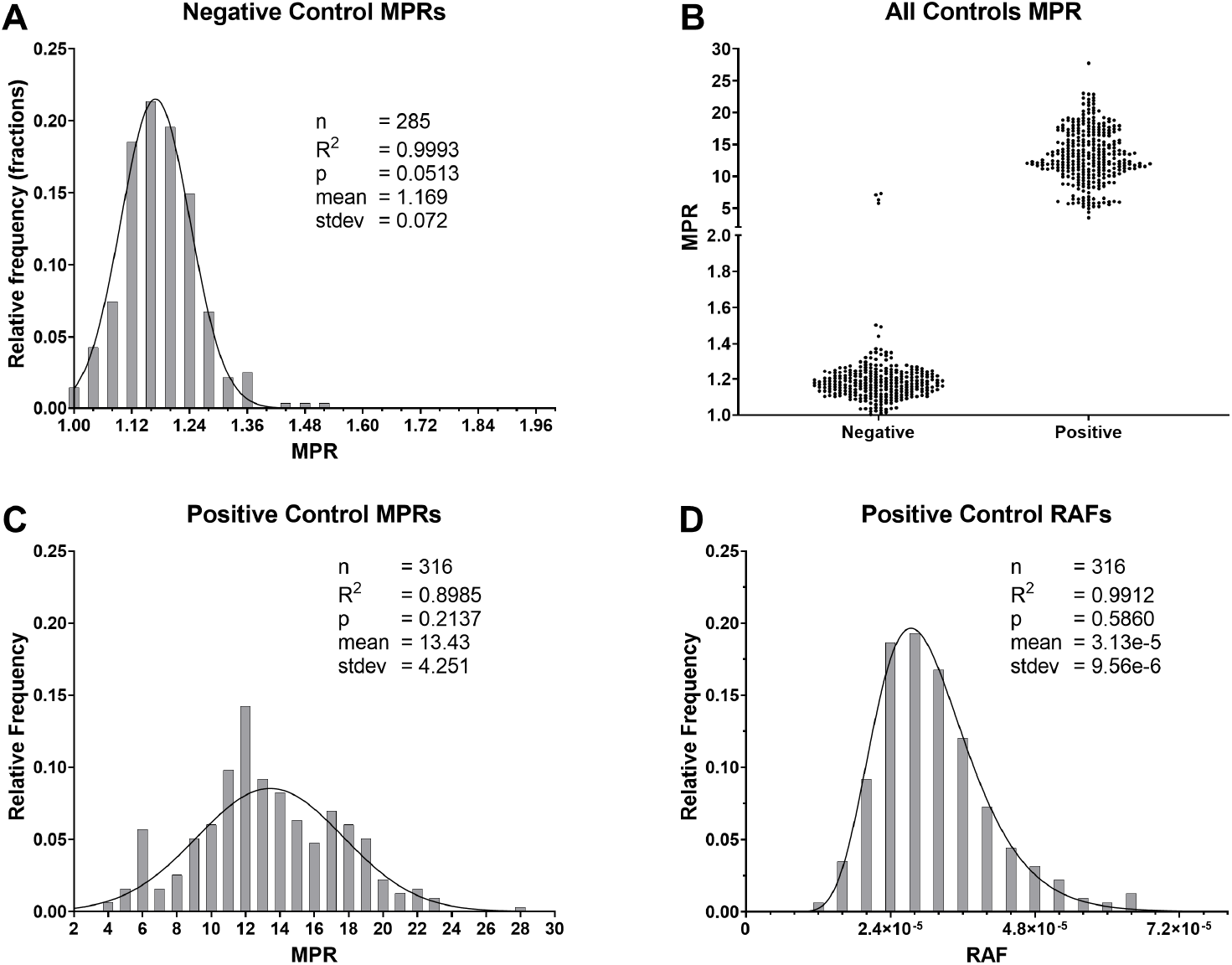
Isolated maxpoint ratios (MPR) for the RT-QuIC control samples. **A.** Negative control MPRs fit a Gaussian distribution and the values are slightly skewed in the positive direction. **B.** Distribution comparisons of the negative and positive controls. **C.** The MPRs of positive controls fit a Gaussian distribution, however, roughly 10% of the variance was unexplained by the model. **D.** A lognormal distribution was observed for the corresponding rate of amyloid formation values of the positive control with an excellent fit for these values.

### T*_MPR_* and Rate of Amyloid Formation Determinations

A conservative threshold (T*_MPR_*) was set at a value of 2.0 for determining RAF with the MPR method. This T*_MPR_* was based on the distributions highlighted in Figure 1. Example graphs showing the output of this method are demonstrated in Supplementary Figure S1. A receiver operating characteristic (ROC) analysis was performed on individual MPR values for which there were tissue-matched, bilaterally sampled ELISA results (Figure 3). Analysis revealed an area under the curve (AUC) of 0.9226 (95% CI: 0.8853–0.9600) meaning that individual MPR values diagnose in line with ELISA 92.26% of the time. Cutoff values were then sorted in descending order by their Youden’s index values (*J*). The top cutoff value suggested a T*_MPR_* of 1.957 which supported our chosen T*_MPR_* value of 2.0. Notably, five individual WTD were represented by RT-QuIC replicates >2.0 but were negative for ELISA. Two of these samples were initially detected as CWD-positive by ELISA in medial retropharyngial lymph nodes (RPLNs) [51], but not in the tissue matched with these RT-QuIC results.

**Figure 3.**
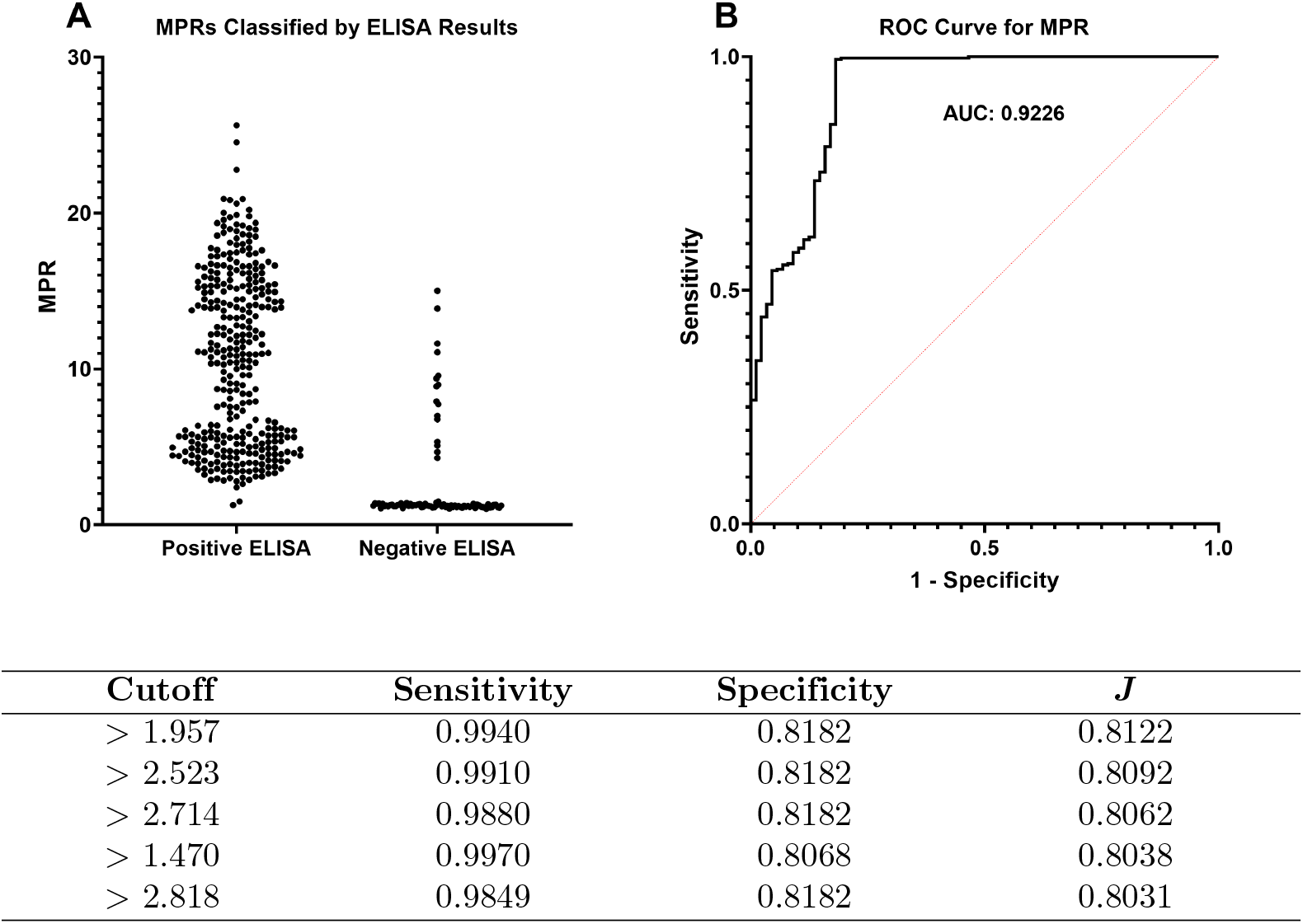
**A.** Maxpoint ratios (MPR) of unknown-CWD-status sample replicates compared to tissue-matched, bilaterally-sampled, enzyme-linked immunosorbent assay (ELISA) results. **B.** Receiver operating characteristic (ROC) curve and its area under the curve (AUC; 0.9226) for individual replicates’ MPR vs ELISA results. The table shows cutoff values with the highest corresponding Youden’s index (*J* = sensitivity + specificity - 1) in descending order. The top cutoff value complimented our chosen (T*_MPR_*) of 2.0 for determining the rate of amyloid formation (RAF).

### Distribution Differences Between Lymph Nodes

When the lower distribution was further divided based on tissue type, parotid (PLN; n=2,594, R^2^: 0.9993; p: 0.0736), submandibular (SMLN; n=217; R^2^: 0.9982; p: 0.1936), and tonsil lymph node (TLN; n=427; R^2^: 0.9919; p: 0.4846) distributions were all still very well explained by the Gaussian regression model (Figure 4A). The p-values for all of the density plots were fairly low, indicating that they all skewed in the positive direction. The multiple comparisons analysis showed that there was no significant difference between any of the tissue-specific distributions (Figure 4B).

**Figure 4.**
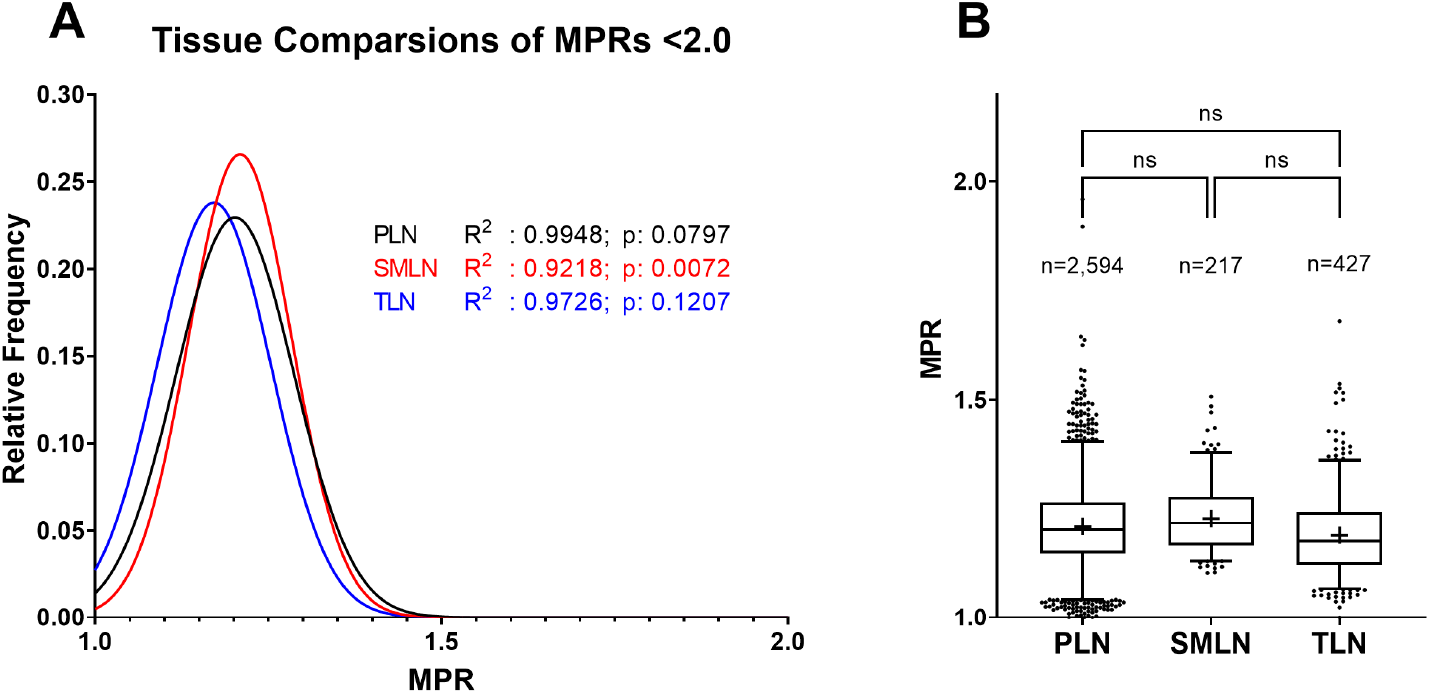
**A.** Comparison of MPRs in the lower distribution separated by tissue type. Medial retropharyngial lymph node (n=17), popliteal lymph node (n=16), and prescapular lymph node (n=30) values were excluded due to insufficient data. Regression analysis shows support for the models regardless of the three tissue types. **B.** The multiple comparisons test found no significance between any of the tissue-specific distributions. Boxes represent the interquartile range, lines are set at the median, “+’s” denote the mean, whiskers are the 2.5–97.5 percentiles, and dots are values lying outside the 2.5–97.5 percentiles.

### Correlation Between MPR and RAF

RAFs were plotted against their corresponding MPRs to determine any relationship between the two (Figure 5). Spearman’s coefficient (r: 0.5449, 95%CI: 0.4943-0.5918) suggests that they are mostly independent of each other. The weak correlation resulted in a poor fit of the linear regression as well (R^2^: 0.2649).

**Figure 5.**
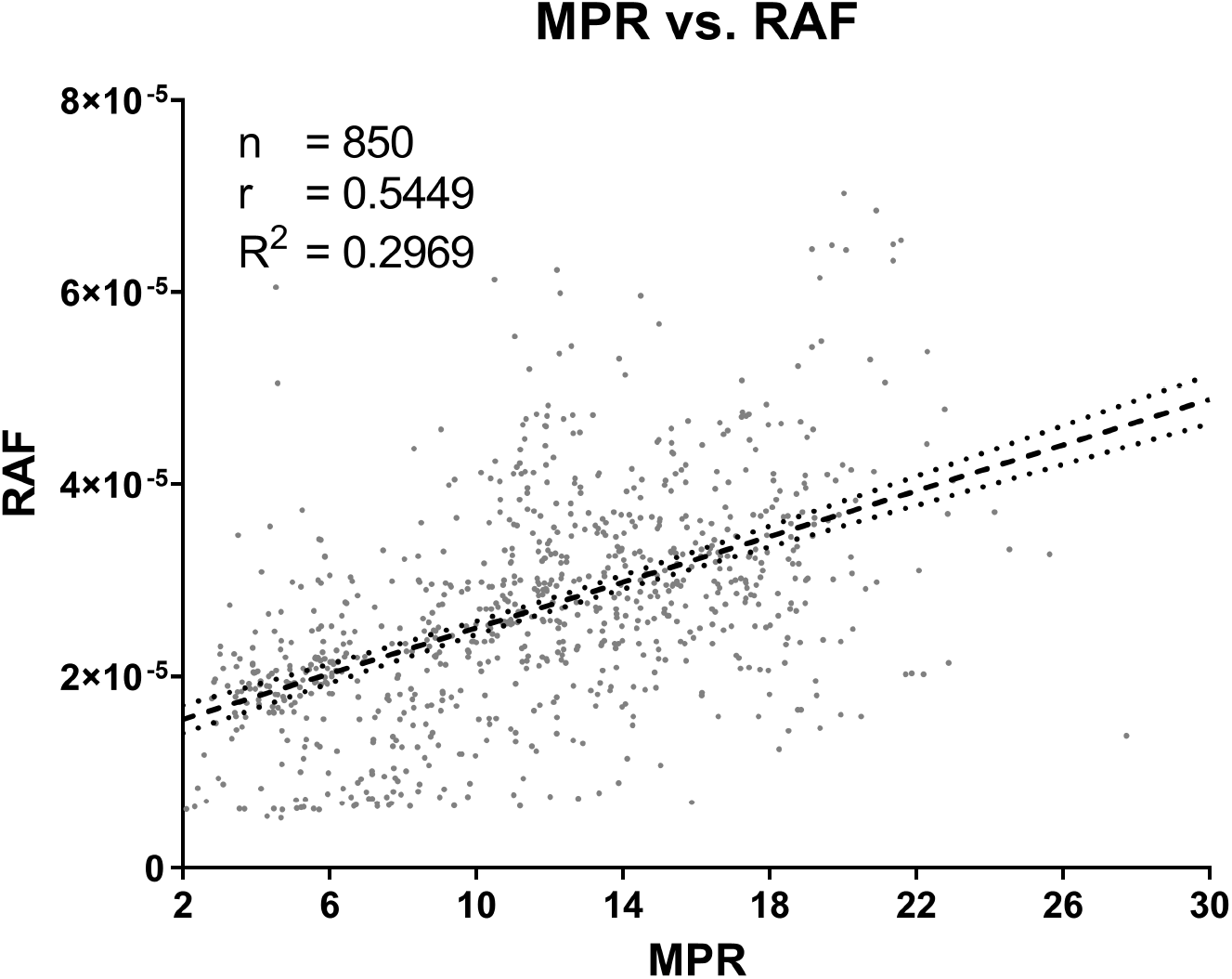
Correlation between maxpoint ratios (MPR) and their corresponding rates of amyloid formation (RAF; i.e., the relationship between signal intensity and reaction efficiency). While there seems to be a slight positive linear relationship, the large amount of scatter and weak correlation suggest that MPR values and their corresponding RAF values are independent of each other.

### Machine and Batch Effects on MPR and RAF

We examined variability of MPRs and RAFs between plate readers and recombinant substrate (recPrP) batches to understand their potential effects on the assay. No significance was detected in negative control MPRs between batches (Figure 6A). However, significant variability was observed in those MPRs when separated based on plate readers (p: <0.0001-0.0129) except for one comparison (1 vs. 4; p: 0.9995; Figure 6B). The only significant batch effects for the positive control MPRs were noticed in batches 1 vs. 3 (p: 0.0004) and 3 vs. 4 (p: 0.0242; Figure 6C). Significance for positive control MPRs between plate readers was detected between readers 1 vs. 3 (p: 0.0007) and 2 vs. 3 (p: <0.0001; Figure 6D). Notably, the first two batches of recPrP (1 & 2) yielded significantly slower RAFs than the subsequent batches (3 & 4) (p: <0.0001; Figure 6E). Significant differences were detected in RAFs between all plate readers (p: <0.0001–0.0321) except 1 vs. 2 (p: 0.9612) and 3 vs. 4 (p: 0.9999; Figure 6F).

**Figure 6.**
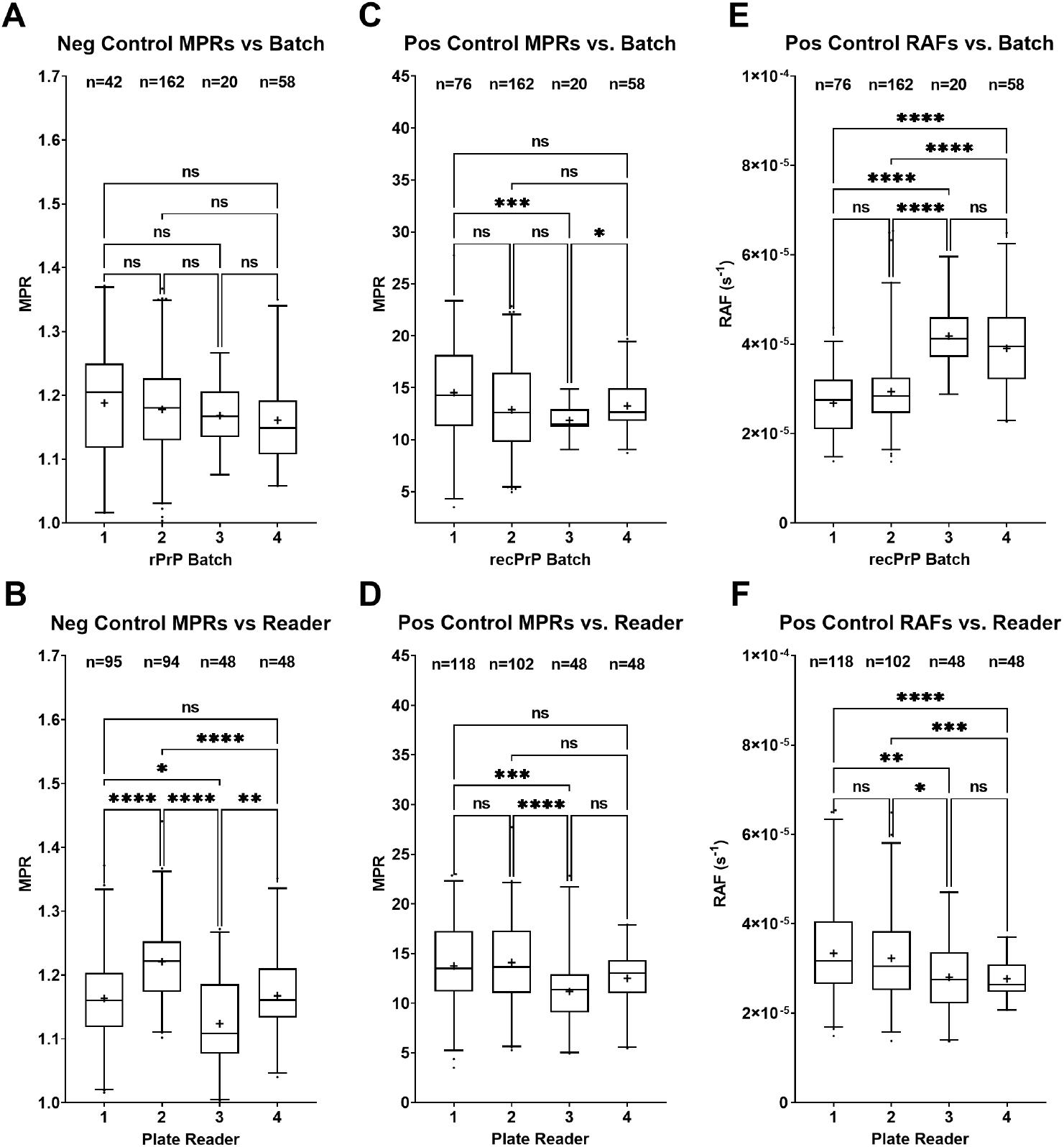
Comparisons of controls depending on the plate reader or recombinant substrate (recPrP) batch used. Boxes represent the interquartile range, lines are set at the median, “+” denotes the mean, and whiskers are the 2.5-97.5 percentiles (****, p<0.0001, ***, p<0.001; **, p<0.01; *, p<0.05; ns, p>0.05). **A.** No significant difference was detected in negative control maxpoint ratios (MPR) between batches. **B.** Significant differences in negative control MPRs between plate readers for every comparison except between plate readers 1 vs. 4. **C.** Significant differences between positive control MPRs in batches 1 vs. 3 and 3 vs. 4. **D.** Significant difference in positive control MPRs between plate readers 1 vs. 3 and 2 vs. 3. **E.** Significance was observed in the amyloid formation rates (RAF) of the positive controls between the first two batches (1 & 2) and the last two batches (1 & 3). **F.** Significant differences in RAFs were observed between all readers except 1 vs 2 and 3 vs. 4.

**Figure 7.**
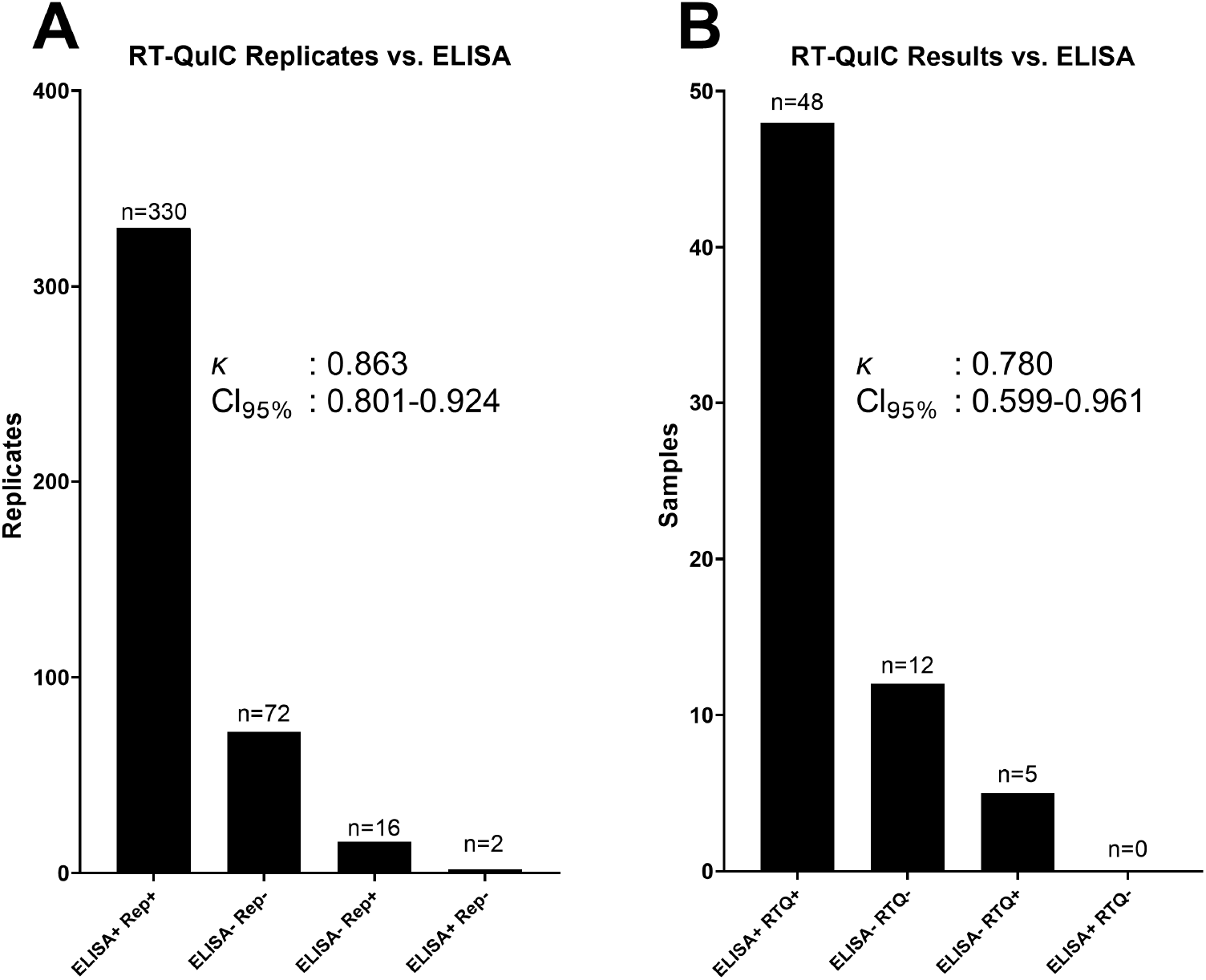
**A.** Replicates that crossed the maxpoint ratio threshold (T*_MPR_*) compared to ELISA results. Of the 420 replicates, 402 agreed with the enzyme-linked immunosorbent assay (ELISA) results. Sixteen replicates crossed T*_MPR_*, but the sample was not detected by ELISA, and two replicates did not cross T*_MPR_*, but the sample was detected by ELISA. **B.** After statistical analysis, 60 samples agreed with the ELISA results, five were positive for real-time quaking-induced conversion (RT-QuIC) but negative for ELISA, and none were negative for RT-QuIC but positive for ELISA.

### Comparison of RT-QuIC to ELISA

Because ELISA is used as the standard screening process for CWD diagnosis, a comparison was necessary to determine the agreement between MPR results and ELISA results. Out of 528 RT-QuIC replicates produced from 64 lymph node samples (with one sample tested twice due to an ambiguous result) for which ELISA data were available, 402 replicates agreed with the ELISA results whereas 18 replicates disagreed (κ: 0.863, 95%CI: 0.801-0.924; Figure 7A). After comparing MPR replicates to their respective plates’ negative controls (such as in Supplementary Figure S1) using ANOVA and a Dunnett’s multiple comparisons test, 60 samples agreed with ELISA, five samples disagreed with an ELISA-negative result, and no samples disagreed with an ELISA-positive result (κ: 0.780, 95%CI: 0.599-0.961); Figure 7B). Of the five that disagreed with the ELISA-negative result, two of those individuals were confirmed CWD-positive in a different tissue.

## Discussion

Currently, there is no standardized, statistical approach for determining prion disease status when utilizing RT-QuIC. The most commonly used method is based on T*_stdev_*, the calculation of which involves acquiring a baseline mean RFU of an entire reaction plate and then determining a T*_stdev_* as a varying integer of standard deviations from that value. RAFs are then defined as the inverse of the time taken to reach the T*_stdev_*; however, the magnitude of these values is typically not considered in diagnostic determinations. Importantly, this method does not take into account initial variation between replicates, samples, or particular experiments. Thus, reliably comparing multiple experiments becomes increasingly complex. Furthermore, meaningful statistics are difficult to perform on the RAFs alone when there is no variance in an ideal negative control (i.e., all replicates have an RAF of zero). In such cases, using a t-test or Mann-Whitney test is not statistically informative because a positive is determined solely based on how many wells cross T*_stdev_*. Just as with RAFs calculated through this approach, those obtained by crossing T*_MPR_* have similar limitations. Therefore, we posit that RAF values, regardless of provenance, are more useful for understanding the seeding kinetics of a given sample than for determining a diagnosis.

Using MPRs instead solves the statistical issues associated with interpreting RT-QuIC data with RAF values alone. Of 3,036 negative sample replicates from over 500 individuals, MPR values were distributed normally (see Figure 1). Further, when the lower distribution was separated based on tissue type, no significant difference was detected (see Figure 4). This means that when comparing an unknown sample to a known negative lymph node sample, researchers can perform a meaningful, parametric analysis that does not rely on arbitrary determinations.

Interestingly, there is seemingly no correlation between individual replicates’ MPR and RAF values indicating that the two measurements are mostly independent of each other (Figure 5). This may be due to the observation that MPRs for a single tissue sample (in this case, the positive control) have a more weakly supported distribution model (Figure 2C), yet the RAF values for the same samples distribute in a highly-supported, Gaussian model (Figure 2D). Further, one would expect MPR values between any given sample to always be similar because the recPrP concentration is identical for every reaction. This of course assumes that all recPrP is consumed during the run and that fibril formation is homogeneous. However, our recPrP batch comparisons (Figure 6) revealed that this assumption of consistency does not necessarily uphold and implies that the reaction kinetics of RT-QuIC are more complex.

The most notable effect on the negative control MPR distribution occurs between plate readers (Figure 6B). Despite running the same program script, there appeared to be some amount of variation due to machine effects. As a note, microplate reader #1 was an earlier model developed by BMG and was not designed specifically for RT-QuIC. This model lacks the plate stabilizers which reduce unintended, sporadic vibrations to the plate. Nevertheless, significant differences were found in the between all readers (except 1 vs. 4). For the positive control MPR distribution, some divergence was found in batch 3, although this may be due to the small sample size (n=20) for that batch (Figure 6C). Additionally, reader 3 showed some significant variance from the other three readers, however, the range of these values was still well within the other three (Figure 6D). RAF values were more susceptible to recPrP batches. Because the rate increased significantly with newer recPrP batches (batches 3 & 4), this effect may be explained by the improved efficiency and fidelity of our purification methods. Furthermore, these machine and batch effects may contribute to some of the variation not explained by the regression models, and further work will need to be done to determine this.

MPR distributions may be influenced by a multitude of variables such as reader settings, recombinant substrate, and tissue type. Therefore, we suggest that T*_MPR_* for RAF calculations be determined by researchers empirically to account for these variables although a tentative T*_MPR_* of 2.0 is conservative given that it is roughly 9.54 standard deviations from the average of the distribution shown herein (Figure 1) and that this cutoff was also supported by ROC analysis where MPRs were compared to tissue-matched ELISA results. Notably, two of the five tissue samples which were negative for ELISA but positive for RT-QuIC came from known CWD-positive animals suggesting that RT-QuIC is likely more sensitive than ELISA. This observation would have artificially skewed the ROC analysis to indicate a lower diagnostic specificity for RT-QuIC than is actually real. Lastly, the gain setting, which amplifies the fluorescent signal to the sensor, is important for acquiring accurate MPR measurements; if a replicate saturates the fluorescent sensor, the actual MPR value is unknown and will skew any statistical analysis. The gain (typically set between 1,000–1,600) can be adjusted appropriately to account for this.

A unique characteristic of this method is that replicates do not necessarily need to cross T*_MPR_* to be determined as significantly different from the negative control using MPR. A clear example of this would be if a sample’s MPR distribution is significantly lower than the distribution of the negative control. However, if a sample’s distribution is <2.0, but is significantly different from the negative control, we suggest re-testing the sample. As a note, ThT fluorescence increases at lower temperatures, so the background measurement should never be taken from the first few minutes of the experiment. The reaction must equilibrate to the experimental temperature to determine an accurate background value. We recommend an equilibration time of at least 45 min.

We propose that using a combination of MPR and RAF allows for a more robust analysis of RT-QuIC data. Across 4,161 replicates, which were run on four separate plate readers and four separate recPrP batches, negative MPRs were distributed normally (see Figure 1). Furthermore, statistical analysis of 64 samples showed that there was excellent agreement with independently secured ELISA results (κ: 0.780) demonstrating that by using a ratio based on a replicate’s independent background fluorescence, RT-QuIC reactions can be normalized between experiments, plate readers, and recPrP batches. Given the expanded usage of RT-QuIC for CWD and other proteopathies and the anticipated approval for regulated testing of CWD in the United States, it is imperative to have a standardized, statistical method that accounts for multiple variables across experiments. We propose that the MPR method presented herein, at least in part, fulfills that qualification.

## Supporting Information

**Figure S1.**
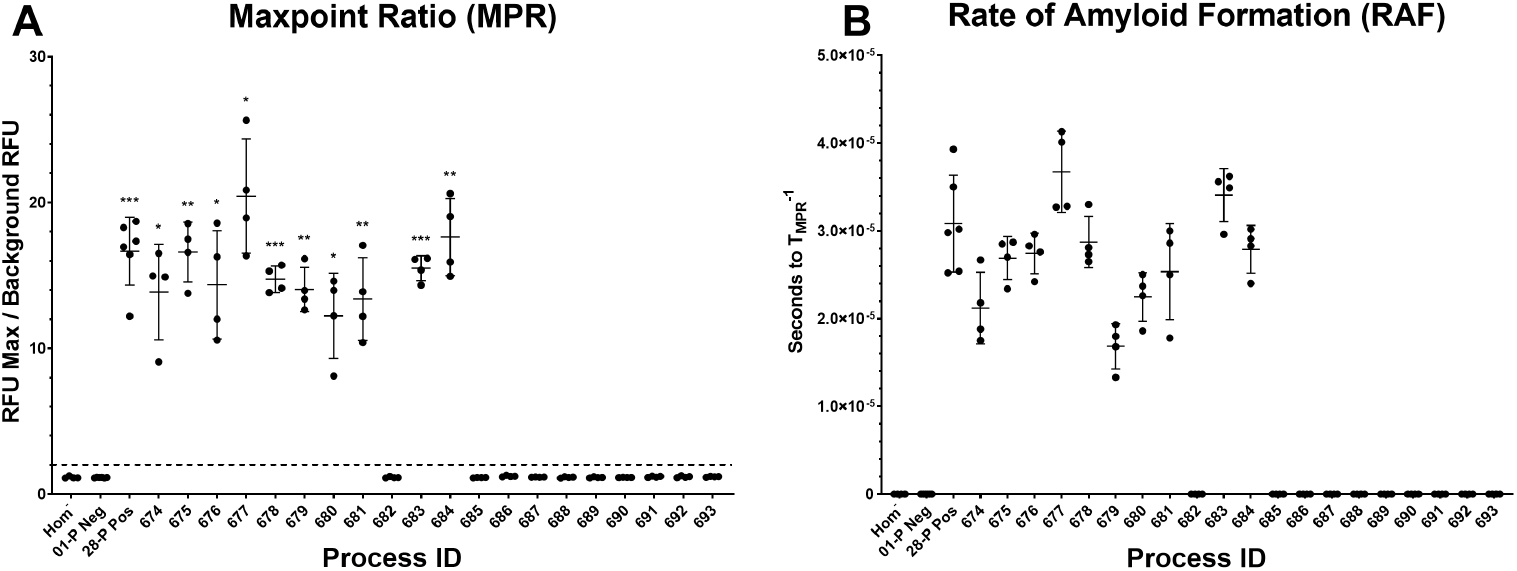
Example graphs highlighting the maxpoint ratio (MPR) and their correlated amyloid formation rates (RAF). **A.** MPRs for each sample are shown with bars at the median extending one standard deviation. The dotted line represents the MPR threshold (T*_MPR_*) for RAF calculation and is set permanently at two. Significance is determined by comparing samples’ MPRs to the negative control MPRs with a Brown-Forsythe and Welch ANOVA and a Dunnett T3 multiple comparisons test (*, p>0.05; **, p>0.01; ***, p>0.001). **B.** RAFs were calculated by taking the reciprocal of seconds needed to pass T*_MPR_*.

## Acknowledgments

We thank Byron Caughey, Andrew Hughson, and Christina Orru of the National Institutes of Health Rocky Mountain Labs for help with establishing and implementing RT-QuIC at the Minnesota Center for Prion Research and Outreach. Fred Schendel, Thomas Douville, and Nicholas Hahn of The University of Minnesota Biotechnology Resource Center were crucial in the production of the recPrP substrate. We thank Kathi Wilson of the Colorado State University Veterinary Diagnostic Laboratory for assisting with ELISA testing. The staff at USDA APHIS Wildlife Services and the Minnesota DNR graciously helped in the collection of WTD samples.

