## Supplementary Figure S1 for "Standardization of Data Analysis for RT-QuIC-based Detection of Chronic Wasting Disease"

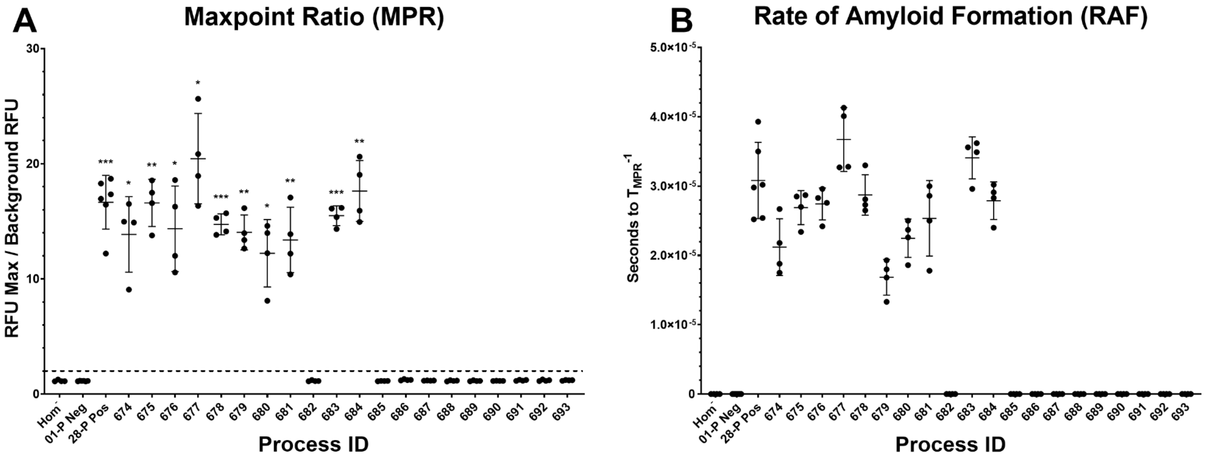


**Supplementary Figure S1** - Example graphs highlighting the maxpoint ratio (MPR) and their correlated amyloid formation rates (RAF). **A.** MPRs for each sample are shown with bars at the median extending one standard deviation. The dotted line represents the MPR threshold ($T_{MPR}$) for RAF calculation and is set permanently at two. Significance is determined by comparing samples' MPRs to the negative control MPRs with a one-way ANOVA and a Dunnett's multiple comparisons test (*, p>0.05; **, p>0.01; ***, p>0.001). (\textbf{b}) RAFs were calculated by taking the reciprocal of seconds needed to pass $T_{MPR}$.
